# From theory to experiment and back again — Challenges in quantifying a trait-based theory of predator-prey dynamics

**DOI:** 10.1101/2021.05.06.442910

**Authors:** K. L. Wootton, Alva Curtsdotter, Tomas Jonsson, H.T. Banks, Riccardo Bommarco, Tomas Roslin, A. N. Laubmeier

## Abstract

Food webs map feeding interactions among species, providing a valuable tool for understanding and predicting community dynamics. Trait-based approaches to food webs are increasingly popular, using e.g. species’ body sizes to parameterize dynamic models. Although partly successful, models based on body size often cannot fully recover observed dynamics, suggesting that size alone is not enough. For example, differences in species’ use of microhabitat or non-consumptive effects of other predators may affect dynamics in ways not captured by body size.

Here, we report on the results of a pre-registered study (Laubmeier et al., 2018) where we developed a dynamic food-web model incorporating body size, microhabitat use, and non-consumptive predator effects and used simulations to optimize the experimental design. Now, after performing the mesocosm experiment to generate empirical time-series of insect herbivore and predator abundance dynamics, we use the inverse method to determine parameter values of the dynamic model. We compare four alternative models with and without microhabitat use and non-consumptive predator effects. The four models achieve similar fits to observed data on herbivore population dynamics, but build on different estimates for the same parameters. Thus, each model predicts substantially different effects of each predator on hypothetical new prey species. These findings highlight the imperative of understanding the mechanisms behind species interactions, and the relationships mediating the effects of traits on trophic interactions. In particular, we believe that increased understanding of the estimates of optimal predator-prey body-size ratios and maximum feeding rates will improve future predictions. In conclusion, our study demonstrates how iterative cycling between theory, data and experiment may be needed to hone current insights into how traits affect food-web dynamics.

## 1 Introduction

Mapping feeding interactions among species in food webs is a crucial first step for understanding how ecological communities function, for gauging the impacts of anthropogenic stress on community structure and stability, and for evaluating how ecosystems might be managed to conserve biodiversity and ecosystem functioning (Thompson et al., 2012). However, to achieve quantitative food-web understanding and predictions, we need a second step of formulating mechanistic models capable of replicating food-web abundance dynamics, and to develop feasible approaches to parameterize such models (e.g. Portalier et al., 2019; Schneider et al., 2012). Only by deriving robust parameter estimates are we then prepared to predict dynamics beyond the range of the existing data, such as what happens when a new species enters the system.

Historically, parameterization of a food-web model required the strength of every trophic link to be independently estimated experimentally, a laborious and often imprecise process (Roslin and Majaneva, 2016). Additionally, some elements, such as non-consumptive interactions among multiple predators, cannot be understood solely from a pairwise predator and prey perspective (Terry et al., 2020). This complexity has made it unwieldy to use dynamic food-web models to map the abundance dynamics of diverse predator-prey assemblages in nature. Recent developments in food-web ecology are now offering a potential cure for this ‘plague of parameters’ (Hudson and Reuman, 2013) through trait-based and, especially, allometric (body-size based) approaches (Yodzis, 1998; Schneider et al., 2012; Boit et al., 2012; Curtsdotter et al., 2019). Such models assume a general relationship between organismal body size and metabolism (Brown et al., 2004; Peters, 1983), and from this infer a relationship between body size and trophic interaction strength (Brose, 2010). Allometric Trophic Network (ATN) models (Otto et al., 2007; Schneider et al., 2012; Berlow et al., 2009) have been formulated based on this idea. They show promising predictions of observed trophic interaction strengths and abundance dynamics of interacting species (Boit et al., 2012; Schneider et al., 2012; Curtsdotter et al., 2019; Jonsson et al., 2018), as well as replication of observed community patterns such as the mass-abundance relationship (Hudson and Reuman, 2013). However, while body size can explain a large portion of observed interaction strengths, in most cases there remains some substantial unexplained variation, which is potentially attributable to other, as yet unmeasured traits (Schneider et al., 2012; Jonsson et al., 2018). Thus, although promising, the general applicability of the ATN modelling approach and the extent to which direct and especially indirect trophic interactions are determined by traits other than body size, remains to be explored.

Among the more successful applications of the ATN model, Schneider et al. (2012, 2014) discuss the potential importance of species’ ‘habitat domain’ (Schmitz, 2007), suggesting that differences in predators’ and prey’s microhabitat use may explain residual variation where the model did not accurately capture the experimental data. Motivated by this, Jonsson et al. (2018) combined the microhabitat use of species with their body size to parameterize an ATN model, thereby successfully predicting experimentally observed population-level interaction strengths when a predator species was alone with its prey (i.e. in the absence of indirect effects from other species). While these and other studies have pointed to the importance of predator and prey habitat use (e.g. Schmitz, 2007; Knop et al., 2014; Staudacher et al., 2018), it has, to our knowledge, never been explicitly incorporated into a dynamic model and parameterized by experimental data. This lack of integration between theory and empirical validation limits our ability to quantify the effect or importance of habitat use.

The trophic interaction modifications (or indirect trait-based effects) observed in treatments with more than two species and pinpointed by Jonsson et al. (2018) are often behaviour-mediated effects of population-level interaction strengths, where changes in the behaviour of a predator and/or its prey is induced by the presence of another species, thereby modifying the per capita interaction strength between the predator and its prey (Terry et al., 2017). Mechanisms include avoidance of intraguild predation and interference among predator species as well as facilitation (Preisser et al., 2007; Kéfi et al., 2012; Sih et al., 1998; Losey and Denno, 1998; Knop et al., 2014). Such interactions are not described by the ATN model, and so the model poorly captures their population-level effects (Terry et al., 2020; Jonsson et al., 2018). Jonsson et al. (2018)’s results strongly suggested that it is a lack of behavior-based non-consumptive interspecific interference effects in the ATN model that is the main cause for its inability to accurately predict trophic interaction strength in more complex webs. Hence, two promising model developments might improve predictions: to consider the spatial niche of species and/or to account for non-consumptive intra-guild interactions.

Here we report on the post-experiment findings of a pre-registered experiment (Laubmeier et al., 2018) aimed at testing the importance of both microhabitat use and non-consumptive predator-predator effects in food-web dynamics. To this purpose, we altered the ATN model to include these factors. We introduced a term for microhabitat use, where predators and prey will encounter each other more frequently the more time they spend in the same area, thereby showing a stronger interaction strength. We also included a term for non-consumptive predator-predator effects, where avoidance of other predators due to the fear of intraguild predation (e.g. Lima, 2002) or interference by other predators decreases predation rate. Finally, to establish whether the effects of microhabitat use and non-consumptive predator-predator effects were sufficiently strong to be observed across a diverse range of predators, we intentionally selected diverse predators covering a range of guilds and feeding modes. To accommodate effects of variation in e.g. feeding mode on optimal predator-prey body-mass ratio for different types of predators, we allowed the value of the optimal predator-prey body mass ratio to vary from predator to predator in the parameter estimation that followed - thereby departing from other studies utilizing the ATN model, such as Schneider et al. (2012) and Jonsson et al. (2018).

To arrive at an optimized design for generating empirical data to inform theoretical models, we (Laubmeier et al., 2018) *a priori* explored the optimal and minimal timing and frequency of experimental sampling to provide sufficient data on population dynamics to enable the use of the inverse method (Chowell, 2017; Banks et al., 2014) for parameter estimation and model testing parameter values of the model. In the current paper, we now perform these steps and test our model against the resulting empirical data, developing a habitat overlap metric in the process.

## 2 The model

We model predator-prey population dynamics in a food web, inferring interaction strengths from body size and microhabitat use. Stronger interactions occur when prey are close to a predator’s optimal prey size, or when predator and prey overlap more in their microhabitat use. To develop our model, we started with the Allometric Trophic Network (ATN) model, which uses body sizes of predator and prey to dictate interaction strengths (Brose, 2010; Otto et al., 2007; Schneider et al., 2012). We then modified the ATN model to include habitat overlap and non-consumptive predator-predator interactions.

Our modified model was published in Laubmeier et al. (2018). Subsequent to publication, we observed that our original formulation for similarity in microhabitat use did not always capture the amount of time predator and prey were in the same location and therefore their likely frequency of interaction. We have, therefore, modified the habitat overlap index to account for this. Below, the entire, updated model is presented and described.

To account for differences in habitat use across species, we divide the habitat into microhabitat zones, quantify the amount of predation that occurs in each microhabitat zone, and sum across all zones. The amount of predation increases with the amount of time spent in the microhabitat (*p*) and decreases with the size of the microhabitat (*A*). When *p*_*i,h*_ is large, species *i* spends more time in habitat *h*, and if *p*_*i,h*_ and *p*_*j,h*_ are both large, we expect species *i* and *j* to encounter one another more often in habitat *h*. Here, we measure *p*_*i,h*_ empirically.

We also introduce a term that describes the decrease in predation by a predator due to non-consumptive effects of other predators. This may include fear of predation, leading to decreased foraging, or physical interference (Preisser et al., 2007; Sih et al., 1998). We propose that the magnitude of this effect depends on the likelihood of predator *j* being intraguild prey to predator *l*, and therefore depends on the expected attack rate of *l* on *j* (*a*_*jl*_). Microhabitat overlap will also affect predator encounters and should therefore affect the magnitude of non-consumptive effects (e.g. Knop et al., 2014). We account for the effects of microhabitat overlap on non-consumptive predator-predator effects in the same way as described above for predator-prey interactions. We sum over the potential attack rates of all species *l* on a single individual of species *j* to account for time spent avoiding or evading species *l* while species *j* is attempting to capture its own prey. The importance of non-consumptive predator-predator effects is described by the scaling constant *t*_0_, where a large value indicates a high penalty to attack rates due to non-consumptive effects. Non-consumptive effects from a conspecific individual may not be distinguishable from non-consumptive effects from another predator species, and so we remove the intraspecific competition term as used in Schneider et al. (2012) from this version of the ATN model and replace it by the more general expression for non-consumptive effects from other predator individuals of any species.

In total, dynamics for the number of individuals *N*_*i*_ of species *i* are therefore given by:

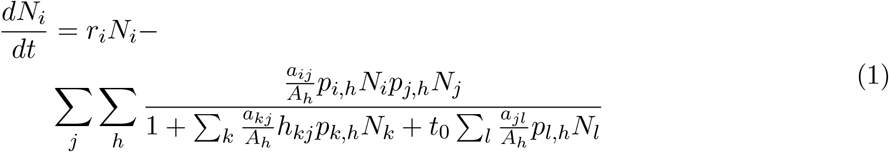

where species *i* increases in proportion to its intrinsic growth rate *r*_*i*_ and decreases due to predation. We assume the intrinsic growth rate (*r*_*i*_) for predators to be zero due to their much longer generation time (a year) compared with the duration of our experiment. The realized per capita attack rate of predator *j* on species *i* in a microhabitat *h* (*α*_*ijh*_) increases with the intrinsic attack rate determined by the predator-prey body-mass ratio (*a*_*ij*_, see below) and decreases with the size of the microhabitat, (*α*_*ijh*_ = *a*_*ij*_*/A*_*h*_), because predator and prey encounter each other less frequently in the larger area. Total predation in a microhabitat increases as the proportion of prey species *i* (*p*_*i,h*_*N*_*i*_) and predator species *j* (*p*_*j,h*_*N*_*j*_), in habitat *h* increases, but decreases dependent on the time predator *j* spends handling prey of the same or other species (*h*_*kj*_), or spends avoiding or interfering with other predators *l*.

As in Schneider et al. (2012), we assume that for species body masses *W*_*i*_ and *W*_*j*_ (corresponding to prey *i* and predator *j*), the allometric parameters (i.e. those dependent on body mass) are given by:

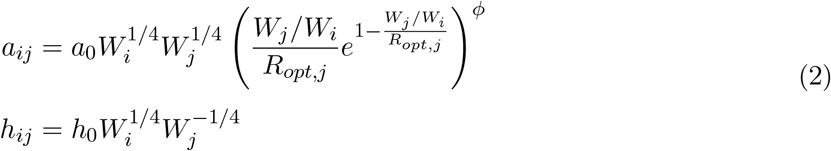

The derivation of allometric parameters is described in Schneider et al. (2012). We note the importance of scaling parameters *a*_0_, *h*_0_, and *R*_*opt,j*_. *a*_0_ scales the frequency of attacks when species encounter one another, with larger values of *a*_0_ indicating more frequent attacks. *h*_0_ scales the time spent handling alternative prey items. Larger values of *h*_0_ indicate more time spent handling prey which results in attacks on a lower portion of the prey population when prey are abundant. *R*_*opt,j*_ indicates the optimal predator-prey body-mass ratio for a successful attack by predator *j*, where *R*_*opt,j*_ = 1 indicates that predator *j* is most successful when attacking prey as large as itself and *R*_*opt,j*_ ≫ 1 indicates that predator *j* is most successful when attacking prey much smaller than itself. Parameter *ϕ* (*ϕ* >= 0) tunes the width of this success curve, with *ϕ* = 0 indicating that attack success is independent of prey size, while the greater the value of *ϕ* the more restricted the attack success around *R*_*opt*_. In contrast to Schneider et al. (2012), we allow the value of *R*_*opt*_ to vary from predator to predator. This is to account for differences in traits not accounted for in the model that may affect predator foraging behavior.

To determine the importance of the terms we introduce — microhabitat overlap and non-trophic predator-predator effects — we compare four variations of the model:

1. the full model (Eq. 1, i.e. with *t*_0_ > 0 and including *p*_*i,h*_ and *A*_*h*_)
2. an intermediate model with habitat use but without predator interference (setting *t*_0_ = 0)
3. an intermediate model with predator interference but without habitat use (removing *p*_*i,h*_ and *A*_*h*_)
4. a minimal model without habitat use or predator interference (removing *p*_*i,h*_ and *A*_*h*_ and setting *t*_0_ = 0)

The different parameters of Eqs 1-2 affect (i) which prey a predator is most likely to consume and (ii) to what extent. As such, the values that we estimate here for *a*_0_, *h*_0_, *R*_*opt*_ etc. for our four model variants will, to some extent, reflect how each model emphasizes the importance of different factors for each interaction. To begin with, a predator with an *R*_*opt*_ value of 1 will interact most strongly with prey of its own size, while a predator with an *R*_*opt*_ of 100 will more effectively consume prey 100 times smaller than itself. Next, if a predator spends all its time on bean plants (*p*_*i,beans*_ = 1) it will have the strongest interaction with prey that also reside predominantly on beans, and have no interaction with prey that are never on beans. Crucially, when the model includes habitat use, it is the combination of both *R*_*opt*_ and *p*_*i,h*_ that dictates trophic interaction strength. For example, consider prey *x* (size = 10 and *p*_*x,beans*_ = 1) and prey *y* (size = 1 and *p*_*y,barley*_ = 1) with predator *i*. If predator *i* (size = 100 and *p*_*i,beans*_ = 1) only interacts with prey *x*, this is already captured entirely by the microhabitat use terms since the predator spends all its time in the same microhabitat zone as prey *x* and never overlaps with prey *y*. In a model accounting for habitat use (as in models 1 and 2), *R*_*opt,i*_, therefore, will likely have an estimated value near 10, since predator *i* is ten times larger than prey *x*. If, however, we do not account for habitat use (as in models 3 and 4), the model needs some other way to capture that predator *i* does not interact with prey *y* in order to optimize the fit to empirical data. In the parameter estimation this could be achieved by a larger value of *R*_*opt,i*_ (moving its optimal prey size further from prey *y*), or a higher value of *ϕ* (narrowing the effective feeding range), thus absorbing differences based on which terms are present in the model and producing good model fit without necessarily reflecting the ‘true value’ of a parameter. This is important to remember when interpreting the results of the model fitting below.

We do not explicitly include non-consumptive mortality in this model. For the aphid (or basal) prey, mortality not due to predation is included in the growth rate term *r*_*i*_, while for predators the experiment is not long enough that we expect mortality other than that due to intraguild predation. Furthermore, without single individual controls, it would be difficult to separate “natural” mortality from that due to predation or cannibalism.

## 3 Methods

### 3.1 The mesocosm experiment

To empirically test and parameterize this model required a study system with rapid growth of the prey population, a range of body sizes of both predators and prey, and distinct habitat zones. With this in mind, and with the benefit of data from a previous experiment (Jonsson et al., 2018), we assembled a six-species terrestrial arthropod community (Laubmeier et al., 2018) (figure 1) dependent on two species of plants; barley (*Hordeum vulgare*) and fava beans (*Vicia faba*).

**Figure 1:**
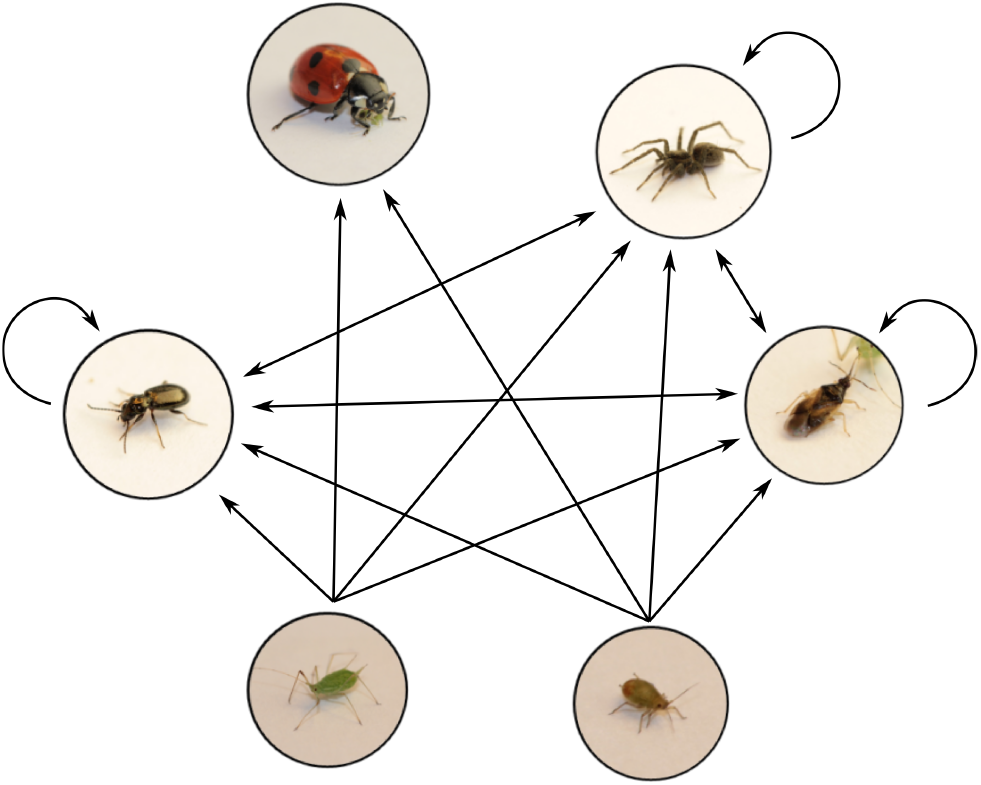
The food web including all possible interactions that we allowed in the model. Species are, from top left: lady beetle (*Coccinella septempunctata*); wolf spiders (*Pardosa* spp.); minute pirate bug (*Orius majusculus*); bird cherry-oat aphid (*Rhopalosiphum padi*); pea aphid (*Acyrthosiphon pisum*); and ground beetle (*Bembidion spp*). Arrows indicate potential feeding interactions which we then parameterized using the inverse method. Arrows point from prey to predator. Double headed arrows indicate that species could potentially eat each other and arrows beginning and ending with the same species indicate cannibalism. We removed all interactions to and from *C. septempunctata* except for *C. septempunctata* preying on aphids, and assumed that the aphids did not consume any predators. This arthropod community was dependent on two species of plants; barley (*Hordeum vulgare*) and fava beans (*Vicia faba*).

As primary consumers we chose one large (*Acyrthosiphon pisum*) and one small (*Rhopalosiphum padi*) species, both aphids. Next, to explore the importance of body mass and microhabitat use in trophic interactions, we chose four predators on these prey, differing in body size and/or habitat preference; one large and one small predominantly foliage-dwelling predator (*Coccinella septempunctata* and *Orius majusculus* respectively), and one large and one small predominantly ground dwelling predator (*Pardosa* spp. and *Bembidion* spp. respectively, where spp. signals the potential inclusion of several congeneric but morphologically indistinguishable species). Each mesocosm contained both barley (*Hordeum vulgare*) (as a host for *R. padi*) and fava beans (*Vicia faba*) (as a host for *A. pisum*), one or both aphid species and zero, one or two predator species. All combinations of predator and prey were replicated six times in a fully factorial design (figure 2). This resulted in 30 predator-prey combinations, plus three control treatments with no predators.

**Figure 2:**
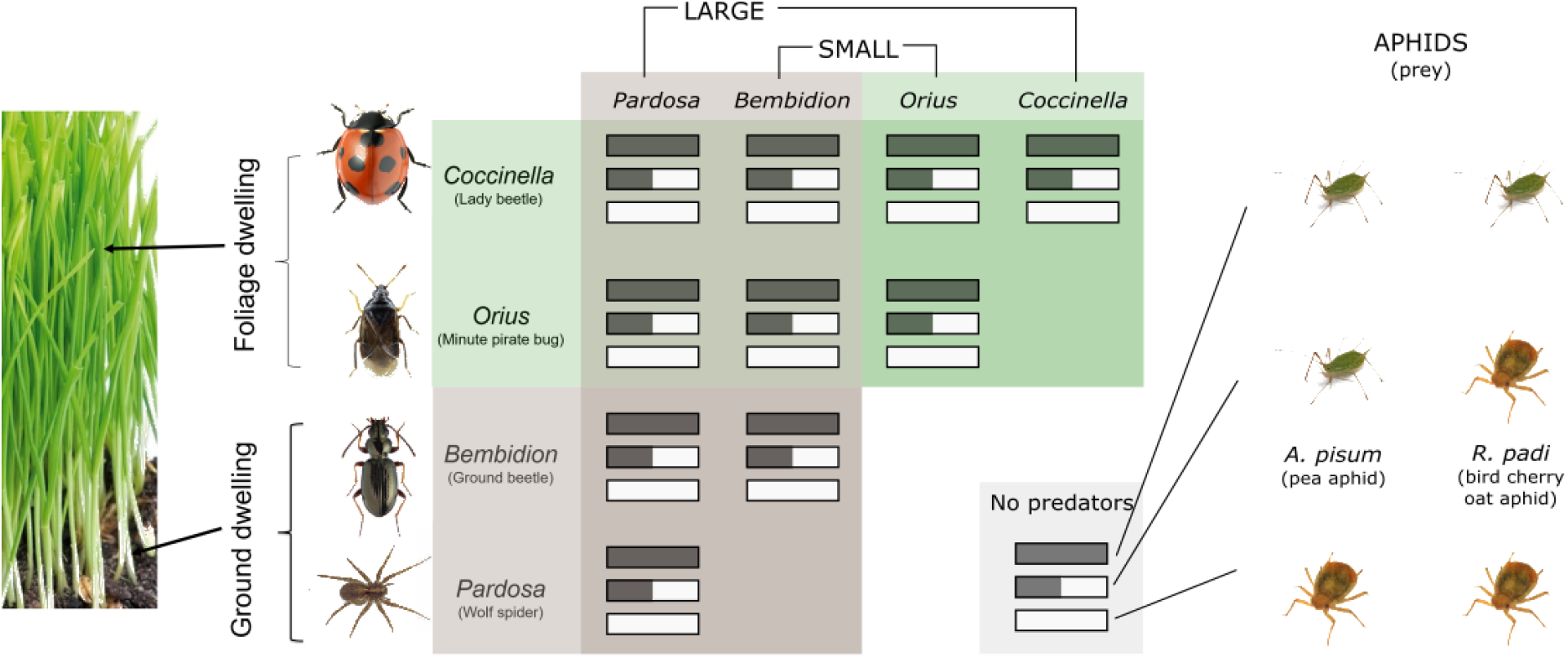
An overview of the predator-prey combinations used in the experiment. Each combination was replicated six times.

Plants were sown in 60×40cm, 20cm deep, plastic containers, sown with two rows of 10 fava bean seedlings and three rows of 15 barley seedlings. A 60cm high mesh cage, with one side resealable to allow aphid counting, was placed on top of each container to prevent insects from entering or escaping the microcosm.

150 wingless adult aphids, placed on Petri dishes, were introduced per microcosm two days before the experiment began. One third of the mesocosms (66 mesocosms) were inoculated with 150 *R. padi* (zero *A. pisum*), one third with 150 *A*.*pisum* (zero *R. padi*), and the final third with 75 *R. padi* and 75 *A. pisum*. Predators were introduced at the beginning of the experiment. The number of predators was determined using a combination of short term (8hr) feeding trials and pilot studies to reach a density where predators would impact the prey, but not eliminate them too quickly. Predator numbers in single-species mesocosms were: *C. septempunctata*: 4 individuals; *O. majusculus*: 40 individuals; *Pardosa*: 20 individuals; *Bembidion*: 40 individuals. Mesocosms with two predators species contained half the number of individuals of each predator species as single predator-species mesocosms, i.e. mesocosms with both *Pardosa* and *Bembidion* together contained 10 *Pardosa* individuals and 20 *Bembidion* individuals

Frequency and timing of aphid counts were determined based on our pre-experimental analyses (Laubmeier et al., 2018). Due to a minor difference in method compared to previous experiments (Jonsson et al., 2018) that made data collection quicker than expected (in-cage rather than destructive sampling), we increased sampling slightly from the minimum determined in Laubmeier et al. (2018). Aphid populations were counted on days 2, 4, 6 and 8. Treatments with *C. septempunctata* were also counted on days 1 and 3, because we realized that *C. septempunctata* decimated aphid populations so rapidly that we would require more data points in order to obtain an estimate of their parameters. Aphids were counted by opening the cage door and carefully counting the number of aphids on each plant.

The proportion of time predators spent in each habitat, *p*_*j,h*_, was measured in single predator mesocosms. The location of each predator was marked on a mesocosm map before beginning to count aphids in these mesocosms. We then categorized these into four areas: walls/roof, ground, beans, and barley. Aphid habitat use was measured by separating aphid counts into each of those categories, but only recorded on days 2 and 6. While the absolute area of beans and barley changed throughout the experiment, we estimated that, on average, the surface area of beans, of barley, and of the ground were roughly equivalent, while the area of the walls and roof was six times larger than each other habitat area (figure 4).

**Figure 3:**
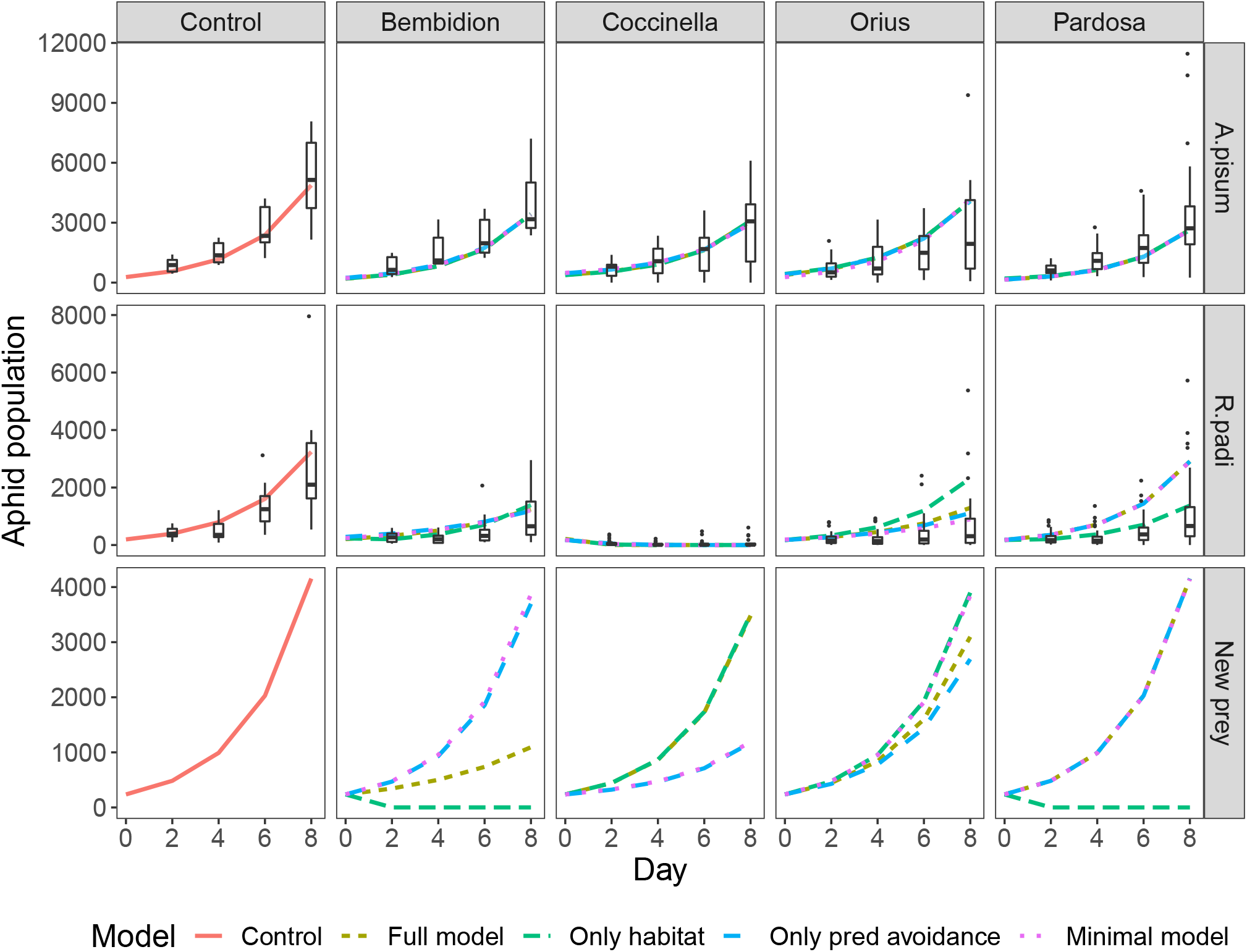
Model predictions of aphid population growth (lines) across time, compared to data of aphid counts per day (boxes) in single-aphid (rows) single-predator (columns) treatments. Lines show predictions of the different models. The final row shows model predictions when fitting each model to a hypothetical new prey species that resides entirely on the ground and has a body size of 1mg.

**Figure 4:**
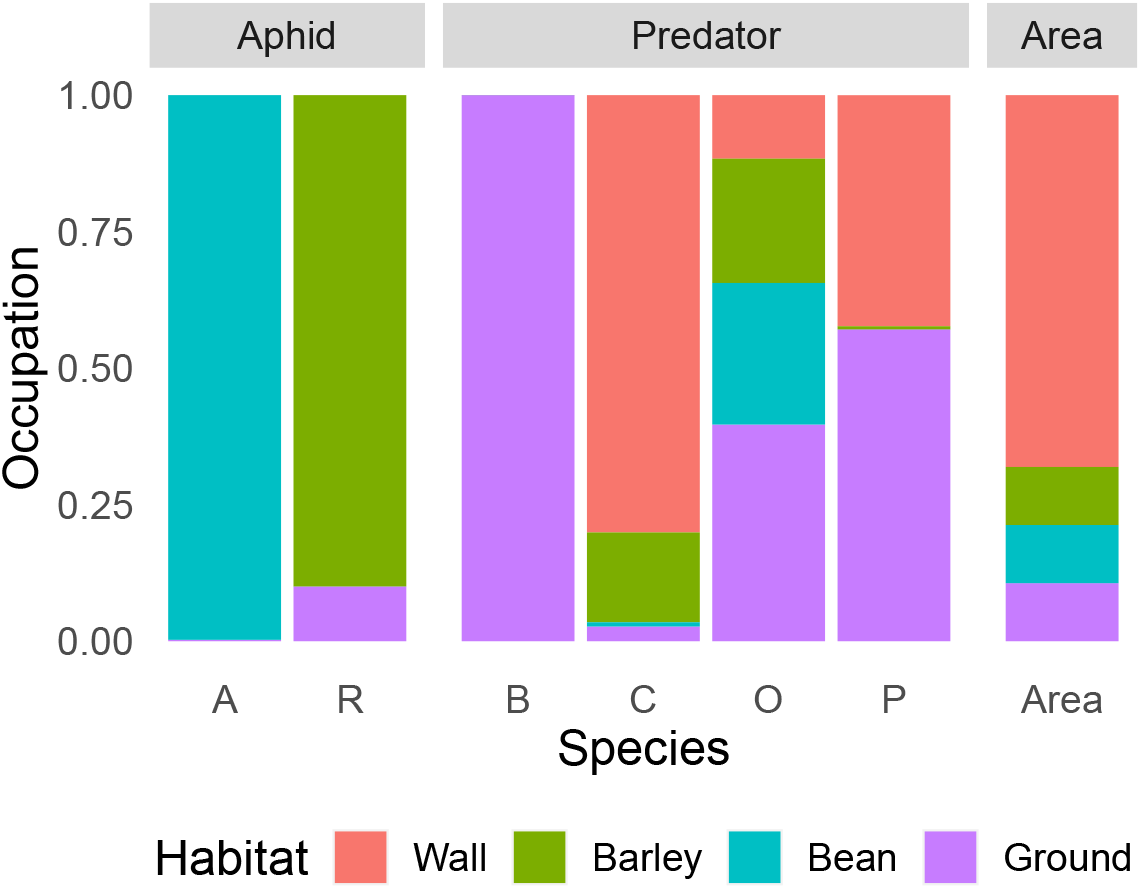
Proportion of time each species spent in each of the four microhabitat zones. The final column shows the relative size of each area. The prey species were A = *A. pisum*, R = *R. padi*, while the predators were B = *Bembidion*, C = *C. septempunctata*, O = *O. majusculus*, P = *Pardosa*.

Predators could only be reliably counted through destructive sampling of the mesocosms, and were therefore only counted on the final day (day 8). After the aphid count, predators were collected by a thorough examination of cage and plant, and sifting through the soil. An additional predator search was repeated the next day to catch any missed in the initial search.

Over the duration of the experiment, we found that *C. septempunctata* could occasionally escape through gaps in the mesh cages. We assume that any *C. septempunctata* missing from cages escaped in this manner, as other predators were never observed consuming *C. septempunctata*. Because this change in the population is not described by our mathematical model, we added replacement individuals to cages where *C. septempunctata* went missing and did not dynamically model the population. Instead, we directly input *C. septempunctata* population densities into the model for other species’ population dynamics. We fixed these densities at constant levels for the duration of the experiment by taking the average value of all observed abundances in each cage. We used averages instead of time-series data due to the uncertainty associated with our observations; it was impossible to know exactly when between observations the individuals went missing from the mesocosm.

The ATN model describes species interaction strengths as a function of species traits (in our case body size and microhabitat use). Because the presence or absence of a food-web link is simply a binary interpretation of interaction strength, the ATN model also predicts the binary food-web structure. However, if a feeding interaction is prohibited due to traits not accounted for in the model, it cannot be expected to correctly predict the absence of such links. As neither body size nor microhabitat could explain why the other predators did not consume *C. septempunctata* (there was microhabitat overlap and predation of similar-sized intraguild prey), we removed feeding interactions between *C. septempunctata* and other predators from the network of potential interactions (figure 1). Similarly, *C. septempunctata* did not consume *Bembidion* or *Pardosa*, for reasons not necessarily explained by microhabitat use or body size (most likely *Bembidion*’s hard cuticle (e.g. Brousseau et al., 2018) and *Pardosa*’s speed), so we removed these interactions.

### 3.2 Model fitting

Using abundance data from our experiment, we parameterized four versions of the ATN model (based on different assumptions for habitat use and predator interference, see below) through the inverse method (Banks et al., 2014; Chowell, 2017). Under this method we first formulated a least squares cost criterion (JLS) (Banks and Tran, 2009; Banks et al., 2014), which describes the deviation of model predictions from empirical observations. Next, by minimizing this cost across all possible parameterizations, we found the best-fitting parameterization. In order to compare the importance of habitat use and predator interference, we repeated this fitting for each of the four models.

To fit the models, we first established a common baseline for aphid growth. Using the data from control treatments, we estimated the intrinsic growth rate (*r*_*i*_) for *A. pisum* and *R. padi*. The rate was distinct for each aphid species, but we assumed the same rate for each aphid species across all aphid treatments (single-species or combined) and replicates. After this, we estimated the remaining model parameters using data from predator-treated mesocosms. There were 10 predator treatments, each with 18 aphid populations; 6 replicates of each of *A. pisum* and *R. padi* in isolation, as well as 6 replicates of *A. pisum* and *R. padi* when in combination with each other.

We simultaneously estimated constants for allometric relationships (*a*_0_, *h*_0_, *R*_*opt,j*_, *ϕ*) and predator interference (*t*_0_) alongside initial aphid abundances. Although model parameters must be the same across all treatments and replicates (but vary among the four models), initial aphid abundances were permitted to vary in every replicate mesocosm. This allowed for differences in population outcomes due to external, potentially stochastic factors, such as variation in plant growth. We constrained estimates of these initial abundances to a range determined by observations from control mesocosms on day 0 of the experiment (ranging from 125 to 775, with a median of 205). Other parameters were unconstrained, except *R*_*opt*_ which was bound between 1 and 1000. This was capped because *R*_*opt*_ values could otherwise reach any upper bound permitted, but in reality increases over 1000 had very little effect on *a*_*ij*_. For the reduced models, we utilized the same values for *r*_*i*_ as in the full model and repeated the process for estimating parameters and initial abundances. To remove predator interference, we set *t*_0_ = 0 and did not estimate that parameter. To remove habitat use, we set *p*_*i,h*_ = 1 and *A*_*h*_ = 1, removing the summation over all *h*.

### 3.3 Model evaluation and prediction

Notably, the above steps utilize the entire data set to derive parameter estimates. Each estimation problem yields a cost criterion (JLS) quantifying model fit, where a lower value of JLS indicates a better model fit. We can also evaluate the performance of each model according to the realism of estimated parameter values and associated processes (e.g. feeding rates), compared to literature or supplemental empirical testing. To summarize, we used a mix of statistical and expert-knowledge model performance criteria, specifically: (i) the JLS cost criterion, (ii) visual fit of predicted vs observed aphid abundance, (iii) realism of parameter values, (iv) realism of ecological processes, and (v) observed vs predicted final predator abundance.

To evaluate performance for these parameter values outside the observed data, we generated predictions for dynamics using an alternative food-web configuration. To demonstrate how the models might compare in their predictions for a new prey species, we used each model, with its resulting parameterization, to predict the population on days 2, 4, 6, and 8 of a hypothetical, entirely ground-dwelling prey species weighing 1mg (slightly larger than *A. pisum*) paired with each of our four predator species.

## 4 Results

All four models gave similar fits to the data (JLS values in table 1 and figure 3), but did so by having different parameter values, particularly *a*_0_ and *R*_*opt,j*_. This is not surprising since both parameters can be expected to absorb differences based on which terms are present in the model, without necessarily reflecting the ‘true value’ of a parameter (see explanation at end of section 2). For example, *a*_0_ scales the frequency of attacks when species encounter each other; a higher value of *a*_0_ signals a higher likelihood of an attack after an encounter and — all else being equal — will lead to a higher predation rate. At the same time, differences in habitat overlap between species decrease encounters, as do non-consumptive predator effects, which — all else being equal — would lead to a lower predation rate. Thus, in models from which the latter two terms are absent, a certain (observed) predation rate can only be achieved by a lower likelihood of attack (i.e. lower *a*_0_) than in models where these terms are present (because here a higher *a*_0_ can be countered by a decrease in attacks due to fewer encounters). In a similar vein, *R*_*opt,j*_ describes the optimal predator-prey body-size ratio for predator *j*. A larger value means predator *j* preferentially attacks prey much smaller than itself, while an *R*_*opt*_ of 1 means that predators preferentially attack prey the same size as themselves. Because we allowed *R*_*opt*_ to vary among predators, we found that a particular predator’s *R*_*opt*_ value could vary substantially among models, thereby compensating for differences in attack rate which were or were not accounted for by habitat use or non-consumptive predator effects in that particular model.

**Table 1:** Parameter values and model fit (JLS) for models with and without habitat use and non-consumptive predator-predator effects. Model 1 = full model with both habitat and non-consumptive effects. Model 2 = only habitat. Model 3 = Only non-consumptive effects. Model 4 = minimal model, neither habitat nor non-consumptive effects. *a*_0_, *h*_0_, *t*_0_ and *ϕ* refer to parameters in Eqs. 1 and 2. *R*_*opt*_.*P, R*_*opt*_.*O, R*_*opt*_.*C* and *R*_*opt*_.*B* refer to the optimal predator-prey body-size ratio for *Pardosa* spp., *O. masculus, C. septempunctata*, and *Bembidion* spp. respectively.

| Model | $a_0$ | $h_0$ | $t_0$ | $\phi$ | $R_{opt.P}$ | $R_{opt.O}$ | $R_{opt.C}$ | $R_{opt.B}$ | JLS |
| --- | --- | --- | --- | --- | --- | --- | --- | --- | --- |
| 1 | 47.37 | 0.014 | 13.66 | 1.41 | 1 | 4 | 1000 | 108 | 1.20e+08 |
| 2 | 7.78 | 0.015 | - | 1.23 | 30 | 997 | 218 | 8 | 1.18e+08 |
| 3 | 0.32 | 0.063 | 9.14 | 1.64 | 1 | 3 | 360 | 389 | 1.22e+08 |
| 4 | 0.39 | 0.057 | - | 1.83 | 1 | 78 | 362 | 377 | 1.38e+08 |

As a result of all this, we saw different parameter values in the different models (Table 1). The different parameter values produced different null-model predictions of feeding rates (i.e. predictions for the scenario where predators and prey used all habitats in proportion to the habitat area; see colored lines in figure 5, here termed ‘*potential* feeding rates’). Despite this, all models produced similar ‘*realized* feeding rates’ of predators on aphids when actual habitat use was taken into account (larger dots at darker vertical lines in figure 5). Consequently, understanding how this is achieved will be useful for deciphering what each model assumes about the foraging behavior of the predators involved.

**Figure 5:**
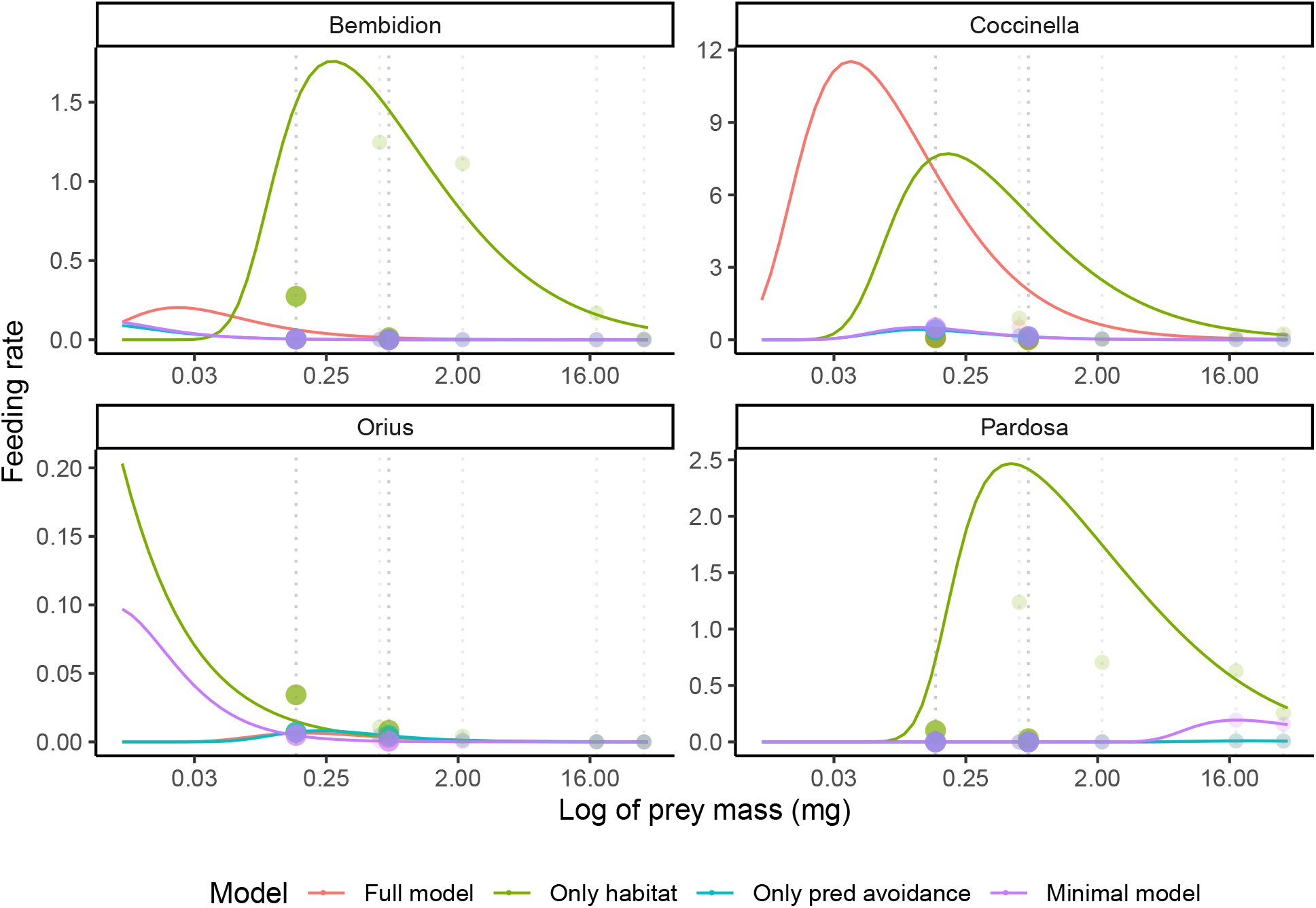
Predictions of each model for each predator population’s feeding rate (y axis) on prey of different body sizes (x axis). Lines show the null expectation or *potential* feeding rate, i.e. predicted feeding rate as a function of prey size if predator and prey used all habitats in proportion to the area of the habitat (no difference in habitat use between predator and prey). This can also be understood as the likelihood that a predator will successfully attack a prey individual *after* they encounter each other, and is clearly much higher for the models that include habitat use. Line color corresponds to different models. Points show *realized* feeding rates of predators on each species in the experiment, based on the prey’s body size (position along the x-axis, denoted by vertical dashed lines) and the actual amount of time predators and prey spend in different habitats. This can be understood as the likelihood that a predator first encounters and then successfully attacks a prey individual. Points are colored according to model, the same as lines. The two aphid species are identified by larger points and darker vertical lines, while predator species (which become intraguild prey to other predators) are shown by smaller points and lighter colored lines. Body sizes were *R. padi* = 0.155mg, *A. pisum* = 0.67mg, *O. majusculus* = 0.58mg, *Bembidion* = 2.15mg, *Pardosa* = 18mg, and *C. septempuntata* = 37mg. Relationships here are shown for predator populations of 20 *Bembidion*, 2 *C. septempuntata*, 20 *O. majusculus* or 10 *Pardosa* individuals with an aphid population of 100 individuals. Despite having a higher *a*_0_ value, full model feeding-rate predictions are lower than the model with only habitat for all species except *C. septempuntata* due to predator-predator non-consumptive effects. Note the varying scales of the y-axis.

As expected, the scaling parameter for attack rate, *a*_0_ was highest in the models with habitat overlap (where encounters were limited), especially the model which also included non-consumptive predator-predator effects (Table 1). *R*_*opt,j*_ values also varied significantly among models, suggesting that this parameter is also absorbing differences based on which terms are present in the model. The optimal predator-prey body-mass ratio for *Pardosa* was equal to one in three out of four models, implying that *Pardosa* is capable of and willing to attack prey of its own size. Such behaviour by *Pardosa* is in line with our own observations made during the experiment, where *Pardosa* individuals readily consumed each other. In models with non-consumptive effects, the optimal prey body size for the small *O. majusculus* was large as well (i.e. small *R*_*opt*_) - a pattern also in line with our observations from the feeding trials, where *O. majusculus* individuals readily consumed *A. pisum* individuals of approximately the same size as themselves. In stark contrast, model 2, which includes habitat use but lacks non-consumptive predator effects, has a value of *R*_*opt,O*_ very close to the upper limit of 1000. In this model with habitat overlap but not non-consumptive predator effects, a small value of *R*_*opt*_ (corresponding to large optimal prey), would result in a much stronger effect of *O. majusculus* on *A. pisum* than we observe in the data. This is for two reasons. First, because *O. majusculus* had relatively high microhabitat overlap with *A. pisum* (figure 4) the model would predict many attacks by *O. majusculus* on *A. pisum* due to many encounters and high *a*_0_. Second, the model predicts that non-consumptive predator effects strongly decrease foraging (and therefore encounters) when *R*_*opt,j*_ is close to one and conspecific individuals become a threat. A model without non-consumptive predator effects, therefore, would not show a decrease in attacks on *A. pisum* due to *O. majusculus* avoiding conspecific potential predators. The high value of *R*_*opt*_ which the model in fact fitted, accounted for the lower-than-otherwise-expected effect of *O. majusculus* on *A. pisum*, by driving down attack rates of *O. majusculus* on all prey. *C. septempuntata* seemed to prefer smaller prey, with *R*_*opt*_ values ranging from 218 to 1000. *R. padi*, which *C. septempuntata* had a very strong effect on, has a body-size ratio with *C. septempuntata* of 242, so this range of *R*_*opt*_ values reflects *C. septempuntata’s* impact on *R*.*padi. Bembidion* also tended to have larger values of *R*_*opt*_, especially when the model did not include habitat use. This may reflect the fact that *Bembidion* has a negligible impact on *A*.*pisum*. When habitat use was included in the model, the negligible effect of *Bembidion* on *A. pisum* was accounted for by the fact that they had very little overlap. Without habitat, the larger value of *R*_*opt*_ predicted a lower interaction strength with *A. pisum*.

Other parameters varied less among models. Handling time was uniformly low across all models, ranging from 0.014 to 0.063. When included in the model, *t*_0_ was high (13.66 and 9.14 with and without habitat respectively), suggesting that non-consumptive predator effects are important in those models. *ϕ* ranged from 1.23 to 1.41, which tells us that the importance of predator-prey body size was roughly consistent across models.

While the models fit very similarly to the dynamics of aphid populations, they gave different predictions of predator populations (figure 6). The full model usually gave the best prediction of final predator population sizes. Omitting only non-consumptive predator-predator effects from the model tended to estimate a stronger decline of predator populations than observed in the data, while omitting both non-consumptive effects and habitat tended to underestimate the decline. The exception here is *Pardosa*, where the full model over-estimated the decline. This is because *Pardosa* had a small *R*_*opt,j*_ value in the full model, making cannibalism a common occurrence.

**Figure 6:**
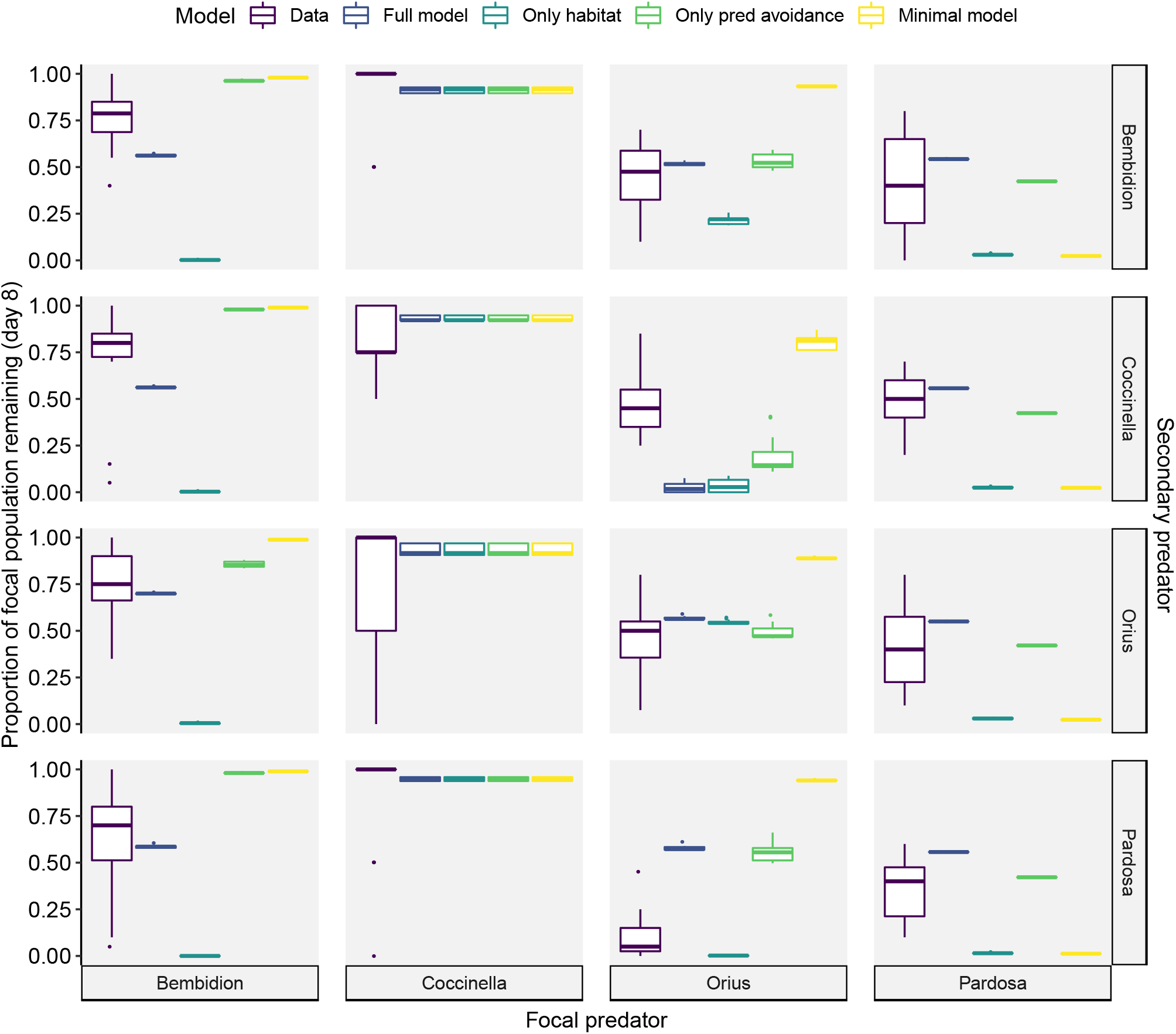
Experimental results (purple boxes) and model predictions for the proportion of each predator population remaining on the final day of the experiment. Data is grouped (facets along the right-hand y-axis) according to the second predator, i.e. the top right panel shows the pro-portion of the *Pardosa* population remaining at the end of the experiment when combined with *Bembidion*. Note that *C. septempuntata* was not modelled dynamically, so all model predictions are the same. Model predictions show different predictions based on different prey treatments and aphid population initial conditions.

Despite the different parameter values estimated for each model, they combined to yield relatively similar *realized* feeding rates (dots in figure 5), especially on the aphids *R. padi* and *A. pisum* (with the exception of *Bembidion* and *O. majusculus* in the habitat-only model). *Potential* feeding rates (lines in figure 5) were higher in the models including habitat overlap, but were (in most cases) dramatically decreased when accounting for the actual habitat use of species. This decrease is shown by the difference between the dots and the lines in figure 5. The few cases where the *realized* feeding rate was larger than the *potential* feeding rate (e.g. the interaction strength of *Bembidion* with itself) occurred when species had similar (or in this case, identical) habitat use, especially in a relatively small habitat area. These *potential* feeding rates are primarily relevant when extrapolating to other species combinations or habitat configurations; when fitting the data from our experiment, it is the *realized* feeding rates, including the effects of habitat overlap and non-consumptive predator-predator effects, that matter. This is why such different models can produce such similar fits to observations.

However, the fact that the different models all produced similar fits to observations, especially of aphid dynamics, does not mean the models are equivalent. When we used the models to make predictions for a new species, each model gave very different predictions (bottom row in figure 3). Using a hypothetical, entirely ground-dwelling prey species slightly larger than *A. pisum*, each model predicted vastly different impacts of the predators on the prey population. Without habitat use, the models predicted that *Bembidion* and *Pardosa* would have no impact on the prey, *C. septempuntata* would be a strong predator, and *O. majusculus* would fall somewhere in between. With habitat, *C. septempuntata* was predicted to have very little impact on the prey (since *C. septempuntata* was almost never on the ground). *Pardosa* and *Bembidion* were predicted to have the strongest effect when habitat, but not non-consumptive effects, was included in the model.

## 5 Discussion

Here, we have reported on the post-experiment results of a pre-registered study aimed at developing a dynamic food-web model, taking into account body size, habitat use, and non-consumptive predator-predator effects. We used the inverse method to determine parameter values of the dynamic model from time-series data from mesocosm experiments. When comparing the fits of four alternative models with or without microhabitat use and non-consumptive predator-predator effects to the dynamics of two aphid species and their predators, we found that objectively (based on the JLS criterion alone) the four models fit equally well and predicted similar aphid population dynamics. Yet, the four models had different values for key parameters, making it difficult to determine which model, if any, could be interpreted as being the best. By having different parameter values, the models ascribed different mechanisms to similar dynamic outcomes. These different mechanisms matter when models are applied beyond the data range, as we showed with vastly different predicted effects of each predator on a hypothetical new prey species across the different models. Below, we examine each finding in turn.

With respect to our model performance criteria, we found that all four models performed relatively similarly. First, the models matched aphid abundances to a comparable extent (JLS range from 1.18 × 10^8^ to 1.38 × 10^8^, table 1), and second, the visual fit was good overall (figure 3). Third, all parameter values were — given our current knowledge — within reasonable ranges (Table 1). Fourth and similarly, processes (e.g. realized feeding rate), also appeared to be within reasonable ranges (figure 5). We found the largest difference between models in our final performance criteria, observed versus predicted final predator abundance. Generally the full model and the model with only predator avoidance gave the closest predictions to the data, but this depended on the predator species (figure 6). Overall, with regards to all five model performance criteria, the models performed well at describing aphid population dynamics for the studied predator-prey combinations. Nonetheless, estimated parameter values varied among models. This variation related especially to attack rate, *a*_0_, and to optimal predator-prey body-size ratio, *R*_*opt,j*_, with more variation in some species than others. This implies that we may ascribe different mechanisms to similar outcomes, whereas the models themselves are far from exchangeable and cannot all be correct at the same time.

That variation in performance between models is so small makes it difficult to confidently assess which model is the most accurate and appropriate. Two criteria point to the minimal model (without habitat use and predator-predator interference) as being the worst: it provided the worst fit (table 1), and it frequently over-estimated both aphid populations (figure 3) and predator populations. (Here, *Pardosa* formed an exception, as the the population of this predator was typically underestimated, figure 6). The full model and the model with non-consumptive predator-predator effects but no habitat use most often yielded the best predictions of the predator populations, while the model with habitat use but no non-consumptive predator-predator effects usually under-estimated the predator population (figure 6). Based on our observations during the experiment, the model with only habitat use also seems to predict the most biased *R*_*opt,j*_ values for both *Pardosa* and *O. majusculus*. We observed that both predator species will readily feed on individuals of a similar size as themselves, yet this model predicts that the optimal prey are 30 or 997 times smaller than *Pardosa* and *O. majusculus* respectively. This would mean that the optimal prey mass of *O. majusculus* is 0.58 micrograms, approximately the mass of a grain of maize pollen (Sheridan, 1982; Porter, 1981). The minimal model also predicted a relatively high *R*_*opt,j*_ value for *O. majusculus*, in contrast to experimental observations. These inaccuracies of the simpler models would hint that the full model, or the model with non-consumptive predator-predator effects but not habitat, is the best model, and that non-trophic predator-predator effects and possibly habitat use are important mechanisms driving trophic dynamics. Each line of evidence is, however, inconclusive, and the variation in performance between models is small. These considerations make it difficult to effectively determine which model is the most accurate and appropriate, and therefore to evaluate the importance (or lack thereof) of habitat use and non-consumptive predator-predator effects.

What our findings suggest is that we are yet to reach the point where we can fit ATN models to data for more diverse species, where traits other than body size also impact interactions, then say with any degree of confidence that this or that model is the correct one and the right one to base predictions upon. Each model points to different mechanisms behind the same outcomes. Models with high *a*_0_ values (those including habitat overlap) ascribe the realized feeding rate to a low frequency of encounters, but predict that when encounters do occur, then an attack is likely. Models with low *a*_0_ values, in contrast, suggest that encounters are common, but that when an encounter occurs, an attack is less likely. Similarly, differences in *R*_*opt,j*_ values attribute different mechanisms to similar predator-prey interaction dynamic outcomes. In both models that include non-consumptive predator-predator interference, the *R*_*opt*_ value of *O. majusculus* is close to the actual body-size ratio between *O. majusculus* and *A. pisum*. This suggests that *A. pisum* is a favoured prey of *O. majusculus*, and that the reason *O. majusculus* does not decimate *A. pisum* is cannibalism. Based on the *R*_*opt,j*_ value of *O. majusculus*, the feeding rate of this species is likely limited by non-consumptive predator-predator effects by conspecifics. In models without this mechanism (non-consumptive predator-predator effects), however, a similarly low value of *R*_*opt*_ would indicate that *O. majusculus* should annihilate *A. pisum* — which it clearly does not. In these models, *O. majusculus* had a larger *R*_*opt*_ value, suggesting that the reason it did not feed on *A. pisum* so strongly was not due to the effects of other predators, but because *A. pisum* was not its optimal prey size. Given that we observed during initial feeding trials that *O. majusculus* would readily feed on *A. pisum*, it seems most likely that non-consumptive predator-predator effects are in fact the limiting mechanism, not incompatible body sizes.

To help differentiate between models and improve prediction accuracy, we will need a closer understanding of which mechanisms are actually at play. Such insight is particularly important for making predictions of novel prey species, as we showed here (figure 3), as well as predictions of changes in existing species. If predator and prey change their habitat use to overlap more or less (as may be caused by loss of habitat for example, e.g. Carroll et al. (2019)), then a model where habitat use is important will have vastly different predictions from one where habitat use is not important. Improving our understanding of the mechanisms underlying food-web dynamics is therefore crucially important before we begin to make predictions. One way to resolve this conundrum, and to enable selection among models with a similar fit to the data, is to explore certain parameters more explicitly, and to estimate their value through supplementary experiments or investigations. Through such added data, we should be able to put limits on the ranges of parameter values, and thereby enable determination of which models are, in fact, most accurate.

In our case, *R*_*opt*_ would be a good candidate. This parameter absorbed a large portion of residual variance in our models, but should be relatively straight-forward to measure in itself (e.g. Brousseau et al., 2018). In our model fitting, we allowed *R*_*opt*_ to vary among predators to account for traits, such as feeding mode, that we had not included in the model. This, however, probably allowed too much flexibility. As a result, the predicted *R*_*opt*_ values varied significantly (in the case of *O. majusculus, R*_*opt*_ covered almost the entire allowed range, from 4 to 997). Feeding trials of each predator with otherwise similar prey covering a range of body sizes should enable a restriction of each predator’s *R*_*opt*_ range. Such additional data would enable these parameters to be fixed or limited, aiding in discerning which models actually are most accurate.

Our study has shown that different model formulations and combinations of parameter values can produce similar outcomes in terms of predicted community dynamics. This implies that success in predicting observed trophic interaction strength and community dynamics of the ATN model so far (e.g. Schneider et al., 2012; Jonsson et al., 2018) must perhaps be interpreted cautiously. More specifically, although it is well established that body size does affect many interactions between species, the generality and exact nature and quantitative form of this relationship needs to be further established. Only then can we rule out that a particular model formulation does not produce the ‘correct results’ (i.e. good fit to the data) for the wrong reasons. For this, the underlying biology of what drives foraging ecology (such as prey preference and feeding behaviour and efficiency) needs to be better understood, especially how it may vary among species and be affected by other traits than body size (e.g. Brousseau et al., 2018). Furthermore, we have shown that it is important to understand model limitations before applying a model to an experimental system, so that some aspect of the system is not more complex than that assumed by the model. In hindsight, we realise that some aspects of our model design might seem overambitious, if the goal were to discriminate between alternative model formulations and estimate parameter values for some universal relationship. More specifically, we explicitly aimed for predator diversity in our experimental design, leading us to include predators like *O. majusculus, Pardosa*, and *C. septempuntata* into the parameterization. As the foraging behavior of some of these predators differs from more traditional ‘grab-and-chew’ predators like *Bembidion*, it is probably too simplistic to describe trophic interaction strengths of all these predators using the same universal relationship and based on one trait (body size) only. On the other hand, our attempt at doing so has revealed the need to develop models that accommodate a diversity of foraging behaviour in predators.

In the current case, we will stop short of these proposed added steps, as the current study was explicitly designed to span the steps reported here (Laubmeier et al., 2018). Our intent is not to arrive at the final solution, but to point to the next step in the iterative process between theory and empirical insight.

### 5.1 Conclusion

In this study, we tested an approach explicitly developed in Laubmeier et al. (2018). Our aim was to arrive at an optimized design for generating empirical data to inform theoretical models. While carefully designed, the real-life implementation of the approach reveals the limitations of the information gained when trying to discriminate between different model versions. This emphasizes the challenges in developing the ATN model approach that need to be tackled. While we were able to arrive at a set of alternative parameter combinations plausible in the light of the data, we were still unable to pinpoint which model is correct without locking down the exact value of one or several further parameters. The importance of doing so was revealed by our exercise of predicting the dynamics of a hypothetical novel species arriving in the system. Depending on which model is true, *Bembidion* and *Pardosa* may have the strongest effect on the novel prey, driving their population size nearly to zero, or they may have no impact and *C. septempuntata* and *O. majusculus* may be the more effective predators. Before we can convert the proposed models to predictive tools, we thus need to do more ground-work and conduct smaller experiments to estimate parameters such as optimal prey body size and attack rate scaling parameters — thereby gaining the resolution to select among multiple models. In conclusion, our study demonstrates how iterative cycling between theory, data and experiment may be needed to hone current insights into how traits affect food-web dynamics.

## 6 Acknowledgements

We owe a huge thank you to the many people who assisted in setting up this experiment, building cages, collecting predators, and counting so many aphids! Especial thanks to Gerard Malsher and Carol Högfeldt for their patient assistance and expertise. Thanks also to the research assistants Joe Bliss, Carly Miranda, Elin Ljunggren, Josefin Farrer-Sundberg, Maylis Moro, Josephine Biro, Lina Wu, Kevin Cestrieres, and Michail Artamonov. Thanks also to Anna Eklöf for valuable feedback on the manuscript.

We acknowledge funding from the Swedish University of Agricultural Sciences, Faculty of Natural Resources and Agricultural Sciences (KLW, TR and RB), Swedish University of Agricultural Sciences, August T. Larsson guest researchers programme (RB), VR 2016-04580 (TR, RB, and TJ) and FORMAS 2016-01168 (TR and RB). AL and HTB were supported by grants AFOSR FA9550-18-1-0457 and NSF DMS-1246991.

